## Supplementary information for "Pressure-driven perfusion system to control, multiplex and recirculate cell culture medium for Organs-on-Chips"

### Supplementary Figures

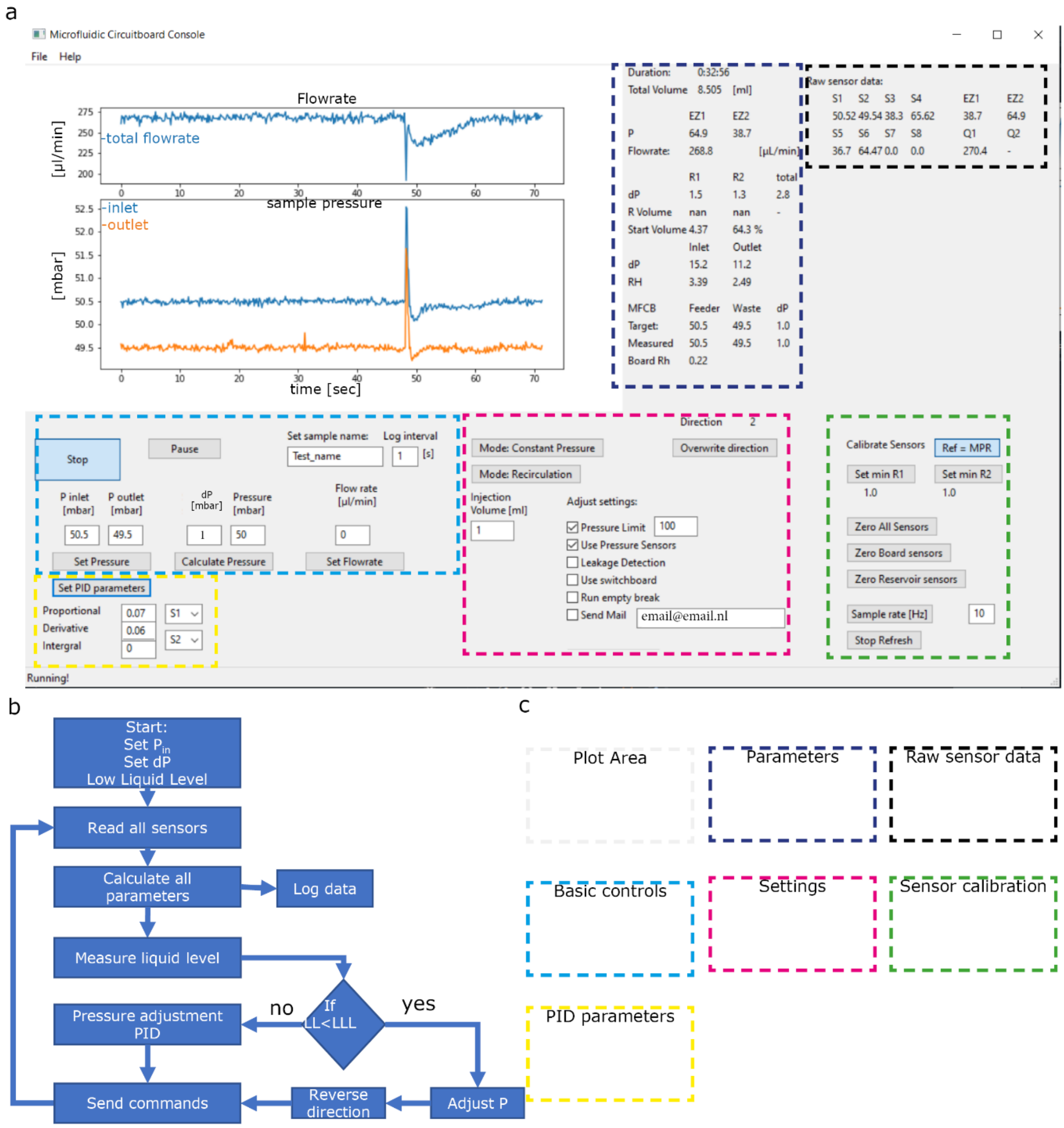


**Fig. s1 Software overview** (a) screen shot of the custom software showing controlling buttons and parameters plots(b) Feedback loop of the software. It starts with reading all sensors and calculate the pressure difference between the inlet and outlet, internal pressure and liquid levels. The PID-controller will adjust the commands to reach the setpoints. (c) Overview of all functions and parameters, explanation in table s1.

| Parameters | Info: |
|---|---|
| Duration | Total experimental time |
| Total Volume | Total displaced volume |
| P | Pressure at controller (filtered) |
| Flowrate | Total flowrate (filtered) |
| dP R1 R2 | Measured P-head (filtered) |
| R volume | Calculate Volume |
| Start volume | Start volume, % measured at present |
| inlet dP | dP inlet Reservoir to Feeder channel |
| outlet dP | dP Waste-channel to outlet Reservoir |
| MFCB | Sample parameters |
| Target Pressure | target difference pressure |
| Measured Pressure | Measured pressure (difference)( filtered) |
| Board Rh | Sample resistance(filtered) |
| Raw sensor data | Unfiltered sensor output |
| **Basic control** | |
| start/stop | Start / stop experiments |
| Pause | No commands, keeps logfile alive |
| P inlet/P outlet | Set target inlet/outlet |
| Set pressure | Sends input to controller |
| Shear/dp/Pressure | set target pressure difference and internal pressure |
| calculate pressure | Sends input to controller |
| Set flowrate | Set flowrate |
| Set sample name | Logfile name |
| log interval | sets log frequency in seconds |
| **PID parameters** | |
| Set PID parameters | Adjust parameters |
| Proportional | P-parameter |
| Derivative | D-parameter |
| Integral | I-parameter |
| Settings | |
| Mode: Constant Pressure / Constant flowrate | Change process variable |
| Mode: Recirculation / Injection | Continuous recirculation, Injection target volume with recirculation |
| injection volume | Target injection |
| overwrite direction | force direction switch |
| Pressure limit | Limits pressure to protect sample |
| Use pressure sensors | Use manufactures PID loop |
| Leakage detection | Switch off when volume is below 85% of start volume |
| Use switchboard | sends 3/2 switch commands |
| Run empty break | If flow>0 break |
| send mail | Sends emails with Situation Reports or leakage/empty break |
| Switch:Medium/Air | Changes algorithm to switch only switch positions |
| **Sensor calibration** | |
| Ref MPR/EZ | Calculate reservoir dP with EZ command or pressure sensor |
| Set min R1/R2 | set low liquid level |
| Zero all sensors | auto zero all sensors |
| Zero board sensors | auto zero board sensors |
| Zero reservoir sensors | auto zero reservoir sensors |
| Sample rate | Change frequency of algorithm |
| Stop refresh | Pauses algorithm |

### Table s2 Channel dimensions FCB

| | Length [m] | Width [m] | Height [m] | $R_h$ |
|---|---|---|---|---|
| Viscosity | | | | 7.90E-04 |
| Feeder channel | 0.2 | 0.0025 | 0.002 | 1.90E+08 |
| Feeder loop | 0.01 | 0.0012 | 0.002 | 4.40E+07 |
| | | | Total | 2.34E+08 |
| Waste channel | 0.2 | 0.0025 | 0.002 | 1.90E+08 |
| waste loop | 0.082 | 0.0035 | 0.002 | 4.33E+07 |
| | | | Total | 2.34E+08 |
| Resistor calculated | 0.010 | 5.00E-04 | 2.50E-04 | 1.77E+10 |
| Total Chip connection | | | | 4.39E+10 |
| Empty microfluidic channel | 0.011 | 5.00E-04 | 5.00E-04 | 4.48E+09 |

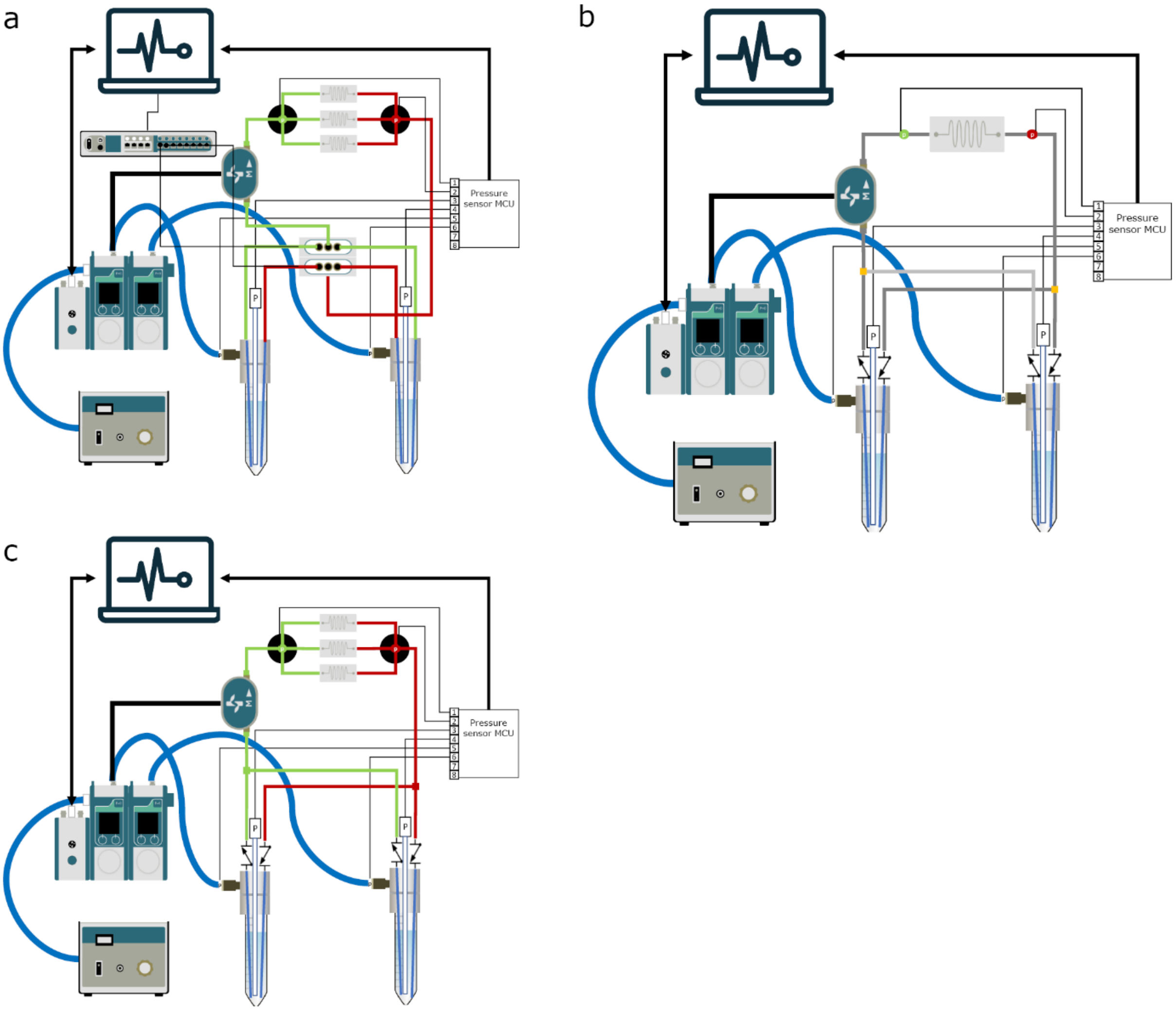


**Fig. s2 Examples of alternative fluidic circuits** (a) recirculation using actively controlled 3/2 valves(Fluigent)(b)Single OoC perfusion,(c) multiplexing using of the shelf manifold splitters.

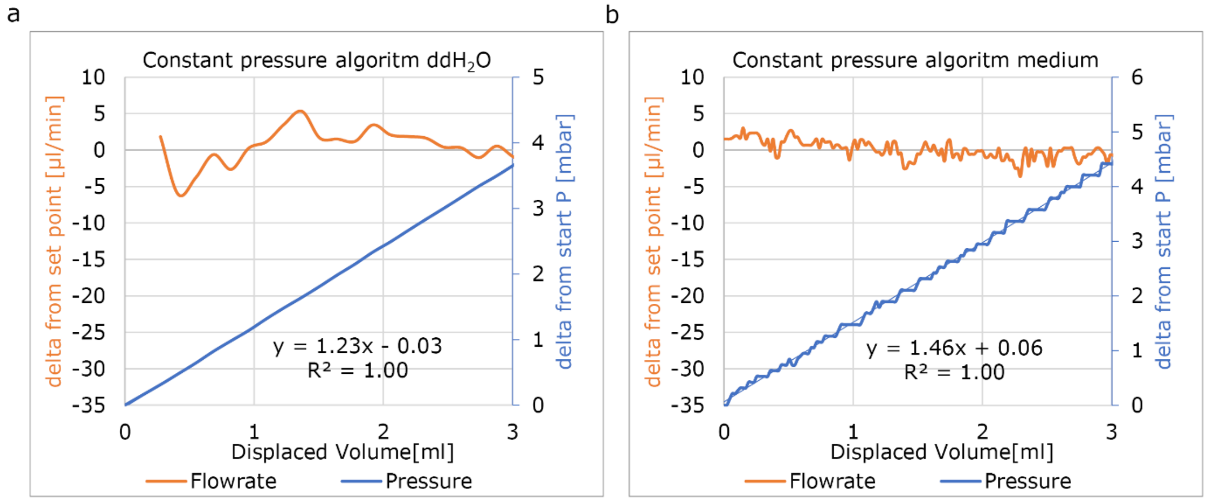


### Fig s3 Typical total pressure increase using the custom PID-controller

(a) Pressure increase per displaced ml of $ddH_20$s. (b) Pressure increase per displaced ml of medium

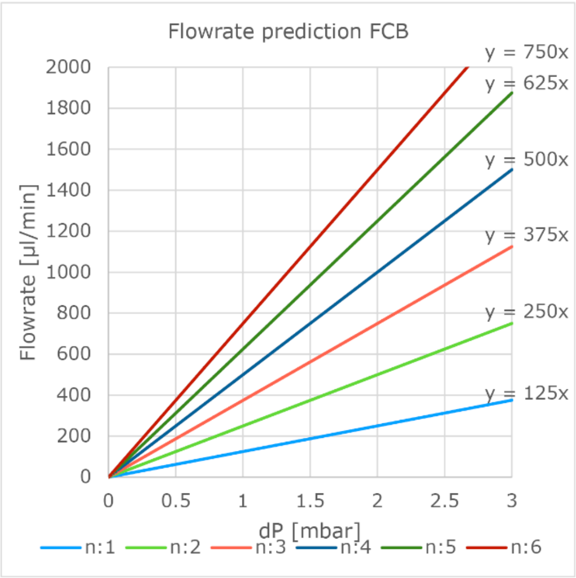


### Fig. s4 Prediction of flowrates

Prediction of multiplexed microfluidic channels(1 to 6) using pressure controlled perfusion. Dimensions listed in **table s2** and EQ 5 and 6 were used to calculate this prediction.
